# Urp1 and Urp2 act redundantly to maintain spine shape in zebrafish larvae

**DOI:** 10.1101/2022.08.09.503396

**Authors:** Anne-Laure Gaillard, Teddy Mohamad, Feng B. Quan, Anne de Cian, Christian Mosiman, Hervé Tostivint, Guillaume Pézeron

**Affiliations:** Molecular Physiology and Adaptation (PhyMA – UMR7221), Muséum National d’Histoire naturelle, CNRS, Paris, France; Structure and Instability of Genomes (String – UMR 7196 – U1154), Muséum National d’Histoire naturelle, CNRS, INSERM, Paris, France; Department of Pediatrics, Section of Developmental Biology, University of Colorado School of Medicine, Anschutz Medical Campus, Aurora, CO, USA

**Keywords:** Zebrafish, Urotensin 2, Urp, spine

## Abstract

Urp1 and Urp2 are two neuropeptides, members of the Urotensin 2 family, that have been recently involved in the control of body axis morphogenesis in zebrafish. They are produced by a population of sensory spinal neurons, called cerebrospinal fluid contacting neurons (CSF-cNs), under the control of signals relying on the Reissner fiber, an extracellular thread bathing in the CSF. Here, we have investigated further the function of Urp1 and Urp2 (Urp1/2) in body axis formation and maintenance. We showed that *urp1;urp2* double mutants develop strong body axis defects during larval growth, revealing the redundancy between the two neuropeptides. These defects were similar to those previously reported in *uts2r3* mutants. We observed that this phenotype is not associated with bone formation defects nor with increased inflammation status but, by using specific inhibitors, we found that the action of Urp1/2 depends on myosin II contraction. Finally, we provide evidence that while the Urp1/2 signaling is functioning during larval growth but is dispensable for embryonic development. Taken together, our results show that Urp1/2 signaling is required in larvae to promote correct vertebral body axis, most likely by regulating muscle tone.

## 1 Introduction

*urotensin 2-related peptide 1* (*urp1*) and *urp2* are two closely related genes encoding neuropeptides. Together with *urotensin 2* (*uts2*) and *urp* (also known as *uts2d* in fishes and *Uts2b* in mammals), they form a multigenic family of neuropeptides that is evolutionarily related to Somatostatin, called the Urotensin 2 family (Tostivint et al., 2006, 2013). These four genes are thought to have arisen through the two events of whole-genome duplication that occurred in the common ancestor of all vertebrates (Tostivint et al., 2013). Characterized first in the Japanese eel (*Anguilla)* (Nobata et al., 2011) and in the zebrafish (Parmentier et al., 2011), *urp1* and *urp2* seem to exist in all fishes but to have been lost in mammals. However, *urp2* has recently also been identified in Xenopus (clawed frogs) (Alejevski et al., 2021). All peptides of the Uts2 family have been suggested to act through a multigenic family of G protein-coupled receptors called Uts2r (also known as Utr or UT). Only one *Uts2r* gene exists in mammals but up to five genes (*uts2r1-5*) have been identified in different species such as the zebrafish and the lizard Anolis (Tostivint et al., 2014).

We have previously reported that in zebrafish, both *urp1* and *urp2* (*urp1/2*) are primarily expressed in the spinal cord in a population of cells called cerebrospinal fluid contacting neurons (CSF-cNs) (Parmentier et al., 2011; Quan et al., 2015). These cells, first described by Kolmer and Agduhr, and also known as KA cells, are GABAergic sensory neurons present all along the spinal cord. They are divided into two subpopulations that have different developmental origins, one located ventrally to the central canal (CSF-cNs’) and another located more dorsally (CSF-cNs’’) (reviewed in (Djenoune and Wyart, 2017; Yang et al., 2020)). We have shown that in zebrafish, *urp1/2* are co-expressed in ventral CSF-cNs both in the embryo and in the adult (Parmentier et al., 2011; Quan et al., 2015).

Lately, *urp1/2* have been suggested to be involved in body axis morphogenesis in zebrafish. First, it was found that the expression of *urp1/2* is reduced in embryos mutant for the cilia motility (ciliary) gene *zmynd10* (Zhang et al., 2018). This mutant presents an embryonic downward curvature of the body axis or curled-down phenotype. This phenotype is a common feature for ciliary mutants (Brand et al., 1996; Jaffe et al., 2016; Kramer-Zucker et al., 2005; Sullivan-Brown et al., 2008) and is thought to be caused by a defect in cerebrospinal fluid flow as a consequence of defective cilia motility (Grimes et al., 2016; Zhang et al., 2018). It has also been reported that overexpressing *urp1* or *urp2* by injection of mRNA or synthetic peptides can partially rescue the curled-down phenotype of *zmynd10* mutants, while it induces the opposite phenotype in wild-type embryos, resulting in a curled-up phenotype. Conversely, in wild-type embryos, morpholinos targeting *urp1* alone or both *urp1* and *urp2* can induce a curled-down phenotype, similar to that of *zmynd10* mutants (Zhang et al., 2018). Urp1/2 peptides were suggested to act through the receptor Uts2r3 which is expressed in dorsal somitic muscles in embryos. Indeed, zebrafish embryo morphants for *uts2r3* appear insensitive to *urp1/2* overexpression (Zhang et al., 2018). Embryos mutant for *uts2r3* do not exhibit any embryonic defect but develop spine deformation during larval growth. Yet, morpholinos targeting *uts2r3* provoke an embryonic curled-down phenotype. The discrepancy between *uts2r3* mutant and morphant phenotypes was suggested to be due to genetic compensation in the mutant (Zhang et al., 2018).

In parallel with this work, other studies shed new light on the mechanisms involved in body axis formation by focusing on the Reissner fiber (RF). This structure, found in most vertebrates, is an extracellular continuous thread, bathing in the CSF in the brain and all along the central canal. This fiber is primarily made of a glycoprotein called subcommissural organ (SCO)-spondin that is produced by the SCO and by the floor plate (Didier et al., 1992; Lehmann and Naumann, 2005; Lichtenfeld et al., 1999; Meiniel et al., 2008; Rodríguez et al., 1998). In zebrafish, the RF starts to assemble during late somitogenesis (20-24 hours post-fertilization [hpf]) and seems to be continuously renewed in a movement from the brain toward the tail (Troutwine et al., 2020). It was shown that several ciliary mutants fail to form the RF, suggesting that functioning cilia are required for normal assembly of the RF (Cantaut-Belarif et al., 2018). Conversely, null mutants for the *scospondin* gene (*sspo)* that have no RF exhibit a curled-down phenotype during embryogenesis, without detectable alteration of cilia nor of CSF flow (Cantaut-Belarif et al., 2018). Altogether, these results suggest that the absence of RF is the cause of the embryonic curled-down phenotype observed in ciliary mutants (Cantaut-Belarif et al., 2018).

More recently, several studies revealed a direct link between the RF and *urp1*/*2* expression. Indeed, similarly to what has been described in the *zmynd10* mutant, the expression of *urp1/2* genes was found to be reduced in *sspo* mutant embryos whereas overexpressing *urp2* appeared sufficient to rescue the curled down phenotype of *sspo* mutants (Cantaut-Belarif et al., 2020; Lu et al., 2020; Rose et al., 2020). In addition, the RF-dependent mechanisms controlling *urp1/2* expression seem to involve the monoamine compounds epinephrine and norepinephrine. Indeed, treatments with these monoamines can restore the expression of *urp1/2* in both *sspo* and *zmynd10* mutants and can ameliorate their phenotypes (Cantaut-Belarif et al., 2020; Lu et al., 2020; Zhang et al., 2018). Moreover, norepinephrine is present in the CSF, in close vicinity of the RF while the adrenergic receptor Adrb2 is expressed in CSF-cNs neighboring cells (Cantaut-Belarif et al., 2020). Finally, morpholino-mediated inhibition of *adrb1* and *adrb2b* genes induces a curled-down phenotype, that can be restored with *urp1* overexpression (Wang et al., 2020).

All these results suggest a model in which the cilia-dependent flow of the CSF is required for the correct formation of the RF. In turn, the RF allows the transport of monoamines, which indirectly signal to the CSF-cNs. These cells secrete the neuropeptides Urp1/2 that act through Uts2r3 on dorsal somitic muscles to promote axis straightening, possibly through the regulation of muscle tone. Of note, we recently reported that *urp2* exists in clawed frogs (Alejevski et al., 2021). In *Xenopus laevis* (African clawed frogs), we found that *urp2* is produced by the CSF-cNs, while *utr4*, the Xenopus counterpart of *uts2r3,* is expressed in dorsal somites during embryogenesis. Furthermore, CRISPR-based inhibition of *utr4* induces an axis defect in Xenopus tadpoles, similar to that of zebrafish *uts2r3* mutant larvae (Alejevski et al., 2021). This suggests that the function of Urp1/2 signaling in body axis is conserved in at least some tetrapods.

Despite this recent progress, several aspects of Urp1/2 functions remain unclear. Since zebrafish mutant for *urp1* have no defect, possibly because of genetic compensation, and *urp2* mutant has not yet been described, the individual contributions of Urp1 and Urp2 to zebrafish development and axis straightness remain unknown. Furthermore, two studies revealed that neuroinflammation is pivotal for axis defect downstream of both cilia and RF defects (Rose et al., 2020; Van Gennip et al., 2018), but this has not yet been addressed in Urp1/2-deficient backgrounds. Also, most mentioned studies focused on the embryonic curled-down phenotype that causes death at early larval stages, thus precluding analysis at older stages. Still, in specific breeding conditions or using hypomorphic mutations, *sspo* mutants could be raised to adulthood: these animals presented strong spine deformations, suggesting that the RF is also important for spine maintenance (Lu et al., 2020; Rose et al., 2020; Troutwine et al., 2020). Whether Urp1/2 are also contributing to these processes during larval growth is not known.

Here, we further investigate the contribution of Urp1 and Urp2 in body axis formation and maintenance. We report the first characterization of *urp2* mutants, which develop a kyphosis that appears in late larvae (3 weeks), and of *urp1;urp2* double mutants, which present strong spine deformations initiating in larvae (1 week). This phenotype is similar to that of the *uts2r3* mutant, showing that Urp1 and Urp2 act redundantly. We show that bone formation and inflammation status are not altered at the onset of phenotype in *urp1;urp2* double mutants. Rather, using myosin II inhibitors, we provide evidence that Urp1/2 signaling acts on muscle contraction, and using conditional *urp2* overexpression, we reveal that Urp1/2 signaling is functional during larval growth. Finally, none of *uts2r3, urp1, urp2* mutants, and *urp1;urp2* double mutants present any embryonic tail-down defect, even as maternal-zygotic mutants. To rule out potential compensation by close family members, we use CRISPR-mediated gene inhibition in *urp1;urp2* double mutants to produce embryos completely deprived of peptides of the Uts2 family. These embryos do not display a curled-down phenotype, showing that Urp1/2 signaling is dispensable during embryonic development. Finally, Urp1/2 are required during larval growth to allow the formation of a straight spine, most likely by regulating muscle tone via their binding to the Uts2r3 receptor, which we show is present in dorsal muscles at the relevant stages.

## 2 Materials and methods

### 2.1 zebrafish husbandry and lines

Zebrafish (*Danio rerio*) were bred according to standard procedures. Embryos were kept in E3 embryo medium until 5 dpf and larvae between 5 and 15 dpf were raised in water with 5g of sea salt and fed *ad libidum* with rotifers. Embryos and larvae were maintained in an incubator at 28.5 °C on 14-10 hr light-dark cycle. From 15 dpf onward, fish were maintained between 26 and 27°C on the same light cycle and fed once a day with artemia and twice with dry food. Wild-type animals were Tübingen obtained from TEFOR-AMAGEN (http://www.celphedia.eu/en/centers/amagen). The *ubi:CRE^ERt2^* line was described previously (Mosimann et al., 2011).

All procedures were approved by the Institutional Ethics Committee Cuvier at the Muséum national d’Histoire naturelle (APAFIS#6945, #19252 and #32413). In accordance with the European Communities Council Directive (2010/63/EU), all efforts were made to minimize the number of animals used and their suffering.

### 2.2 CRISPR/Cas9 RNA guide design and injection

To produce deletion alleles for *urp1, urp2* and *uts2r3*, gRNAs were designed using CRISPOR (http://crispor.tefor.net/) (Haeussler et al., 2016) and produced *in vitro* as previously described (Auer et al., 2014). Ribonucleoprotein complexes were prepared individually by mixing gRNAs with homemade Cas9 protein (gRNA at 80ng/µl and Cas9 at 6µM) and incubated at 37°C for 10 min, to allow the Cas9 protein to bind to the gRNA, and then stored on ice. For each line, two gRNA-Cas9 complexes were mixed and injected (2 nl) into 1-2 cell-stage embryos. To establish lines, injected embryos were raised to adulthood. These founders were crossed against wild-type fish to screen for their ability to transmit a deleted allele by genotyping their progeny. F1 from identified founders were raised to establish the different mutant lines.

For *uts2a*, *uts2b*, *urp* and *pkd2* CRISPR-mediated inhibition, guides were designed using Integrated DNA Technologies design tool (IDT, https://eu.idtdna.com/pages) and crRNA ordered from IDT. According to manufacturer instructions, gRNAs were obtained by producing duplexes with tracrRNA. Ribonucleoprotein complexes were prepared individually by mixing gRNAs with homemade Cas9 protein (gRNA at 13.3µM and Cas9 at 10µM) and incubated at 37°C for 10 min and then stored on ice. Independent gRNA-Cas9 complexes were mixed and injected (2 nl) into 1-2 cell-stage embryos.

All gRNA sequences are indicated in the key resources table. Sequences of oligo for genotyping are indicated in table S1.

In the experiment with *uts2a*, *uts2b* and *urp* CRISPR-mediated inhibition, the efficiency of mutagenesis was assayed by sequencing. A pool of 10 injected embryos at 1 dpf was used to extract genomic DNA and the three targeted region were amplified by PCR (oligo sequences in table S1). Amplicons were cloned into pGEM-T (Promega) using manufacturer instructions. For each gene, plasmids from 10 clones were Sanger sequenced (http://eurofinsgenomics.eu).

### 2.3 Alizarin and calcein staining

For alizarin staining, fish were euthanized and fixed in 4% paraformaldehyde in PBS (4% PFA), for 2-3 days at 4°C. Animals were then rinsed several times in PBS and the skin and all internal organs were removed. Samples were rinsed in water and treated with a solution of borax (5%, w/v, Sigma B9876) overnight at room temperature (RT). Samples were rinsed in water and then in KOH 1% and stained in a solution of 0.015% alizarin red S (Sigma A5533) in KOH 1% at 4°C for 3-4 days. Then, to remove and clear soft tissues, samples were treated with a solution of trypsin 1% (Sigma T4799) and borax 2% for 24 h at RT. Finally, bone preparations were extensively rinsed in PBS and transferred in glycerol 80%.

For calcein staining, live larvae were simply transferred into a solution of 0.02% of calcein (Sigma C0875) in water (pH adjusted to 7.5 with NaOH). Animals were kept in this solution for 30 min and then transferred to clean water to remove unbound calcein. Finally, fish were anesthetized (0.016 % MS222 – Sigma A5040) and imaged.

### 2.4 Real time quantitative RT-PCR

Total RNA samples were obtained from dissected tissues from adults or from a pool of embryos or larvae using RNAble solution (Eurobio) and a TissueLyser II system (Qiagen). Total RNA samples were treated with DNAse I (Roche) to remove contamination of genomic DNA and then purified with phenol/chloroform extraction and sample quality was assayed by electrophoresis on 1% agarose gel. cDNAs were obtained using Goscript reverse transcriptase (Promega), with random primers and 2 µg of RNA for each sample. RT reaction was diluted 20-fold for real-time PCR. Quantifications were performed by real-time PCR with specific primer pairs and using PowerUp SYBR green (Applied Biosystem) on a QuantStudio™ 6 Flex Real-Time PCR System (Applied Biosystem). For all RT-qPCR, housekeeping genes were *lsm12b* and *mob4* (Hu et al., 2016). All primer sequences are indicated in table S1.

### 2.5 Production of Tol2 constructs and injection

The *hsp:urp2* tol2 transgenesis vector was already described (Quan et al., 2021). The *ubi:urp2* and *ubi:STOP^lox^-urp2* tol2 transgenesis vectors were produced using plasmids from the tol2kit (Kwan et al., 2007) and Gateway recombination system (Life Technologies). To produce the entry plasmid *p5E-ubi:STOP^lox^* (*pCM351*), the *loxP*-flanked *STOP* cassette from *pDH083* (Hesselson et al., 2009) was PCR-amplified with primers starting at the 5’ end of the *loxP* sequences with added BglII restriction sites, cut with BglII, ligated into BamHI-linearized *pENTR5’_ubi* (*pCM206*) (Mosimann et al., 2011), and the resulting clones verified by restriction digest and sequencing. Then, *p5Eubi:urp2 (Ref)* or p5E*-ubi:STOP^lox^*, pME-*urp2* (Quan et al., 2021) and p3E-polyA were recombined into the pDestTol2CG2 destination vector. For transgenesis, solutions containing transgenesis plasmids (30ng/µl) and tol2 transposase mRNA (25 ng/µl) were prepared and injected (2 nl) into 1 or 2 cell-stage embryos.

### 2.6 Blebbistatin and BTS treatment

Blebbistatin (Bleb, TOCRIS 1760/10) and BTS (TOCRIS 1870/10) stock solutions were prepared by resuspending powder into DMSO at 100 mM. Solutions were aliquoted and stored at -20°C. To block muscle contraction, Bleb and BTS were diluted in E3 embryo medium at 50µM and 200µM respectively. Those concentrations were chosen because they completely blocked embryo movement (both spontaneous and touch-evoked coiling) without inducing death.

### 2.7 4-Hydroxytamoxifen treatment

To induce overexpression of *urp2* in larvae, transgenic *Tg(ubi:cre^ERt2^)* fish were crossed to wild-type and eggs were injected with the *ubi:STOP^lox^-urp2* plasmid at 1-2 cell-stage. Embryos were raised up to 10 or 20 dpf, at which point the STOP cassette was deleted by activation of the CRE with 4-hydroxytamoxifen. To do so, fish were bathed in system water with 4-hydroxytamoxifen (Sigma H7904) at 7.5 µM (in DMSO) in the dark for 1h on three consecutive days.

### 2.8 Image acquisition

Images of embryos and larvae were acquired with an Infinity3 Teledyne-Lumenera camera on an SZX12 Olympus stereomicroscope, or with a QImaging Retiga-SRV camera on a MZ16F Leica stereomicroscope for fluorescent images. Adults fish were imaged with a Panasonic DMC-FZ18 camera. Images were then processed using ImageJ (Schneider et al., 2012). Images were adjusted for brightness, contrast and color balance but no non-linear adjustment was applied.

### 2.9 Statistics and figure preparation

Quantitative data were analyzed using R software (R Core Team, 2022) and the tidyverse package (Wickham et al., 2019). Plots were produced using the ggplot2 package (Wickham, 2016). Figures were prepared using Scribus software (www.scribus.net/).

## 3 Results

### 3.1 Generation of mutant lines for *urp1*, *urp2* and *uts2r3*

To address the function of *urp1* and *urp2* in zebrafish, we used CRISPR-Cas9-mediated genome editing to produce mutant alleles. The *urp1* and *urp2* genes share the same genomic organization, comprising five exons among which the last one encodes most of the secreted mature peptide (Fig. 1A). Thus, for each gene, we aimed at deleting a portion of the fifth exon corresponding to the end of the coding sequence using two guide-RNAs (gRNA) targeting each side of this region (Fig. 1A). We isolated a deletion allele for each gene, most likely to be null (Fig. 1B-C). Because Urp1 and Urp2 have been suggested to act through the Urotensin 2 receptor Uts2r3, we also produced a deletion allele of the gene encoding this receptor. This gene is made of only one coding exon and we used two gRNAs to delete a part of it (299 pb out of 1161pb). The resulting allele encodes a protein made only of the forty-first amino acids (aa), as compared to 386 aa for the wild-type protein, and is predicted to be null (Fig. 1D).

**Figure 1.**
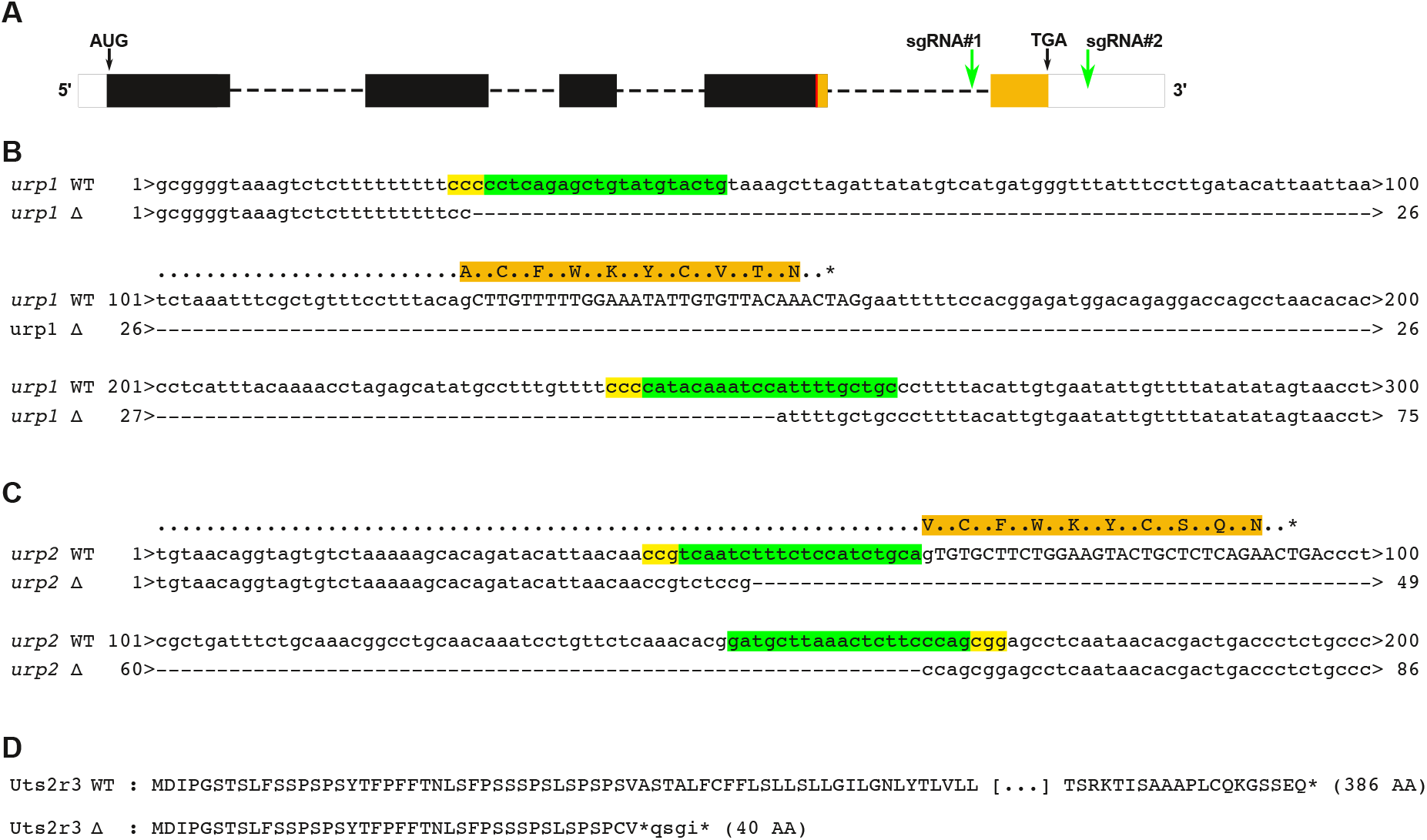
Generation of mutant lines for *urp1*, *urp2* and *uts2r3*. (A). Schematic representation of the genomic organization of *urp1* and *urp2*. Dashed lines indicate intron; white box, 5’ and 3’ UTR; and plain boxes, the coding sequences. The region corresponding to the mature peptide (Urp1 or Urp2) is indicated in orange and is mostly encoded by the fifth exon. In red, the position of the cleavage site. The position of the guide RNAs used to delete the coding region of the fifth exon are also indicated (sgRNA#1, #2, green arrows). (B – C). Sequences from wild-type (WT) and deleted allele (Δ) at the level of the deletion for *urp1* (B) and *urp2* (C). The deletion sizes are 225 pb for *urp1* and 114 pb for *urp2*. The protein sequence is indicated above the wild-type sequence. In green, the sequences of guide RNA with the PAM (NGG) in yellow. (D). Predicted protein sequence for the deletion allele obtained for *uts2r3,* compared to the wild-type sequence.

### 3.2 Urp1 and Urp2 act redundantly for correct spine shape

Zebrafish homozygous mutant for *urp1* developed normally and did not appear different from heterozygous siblings at any stage (not shown), consistent with what was previously reported (Zhang et al., 2018). By contrast, while they initially also developed normally, *urp2* mutants showed a deformation of the spine axis from three weeks (∼8.8 mm Standard length [SL]) onward. This deformation first appeared as a subtle downward bending of the head and progressively became exacerbated in adults, at which stage fish exhibited a strong kyphosis (Fig. 2A-B). Because *urp1* and *urp2* are mostly expressed in the same cells and encode for highly similar peptides (Parmentier et al., 2011; Quan et al., 2015), these observations indicate that *urp1* and *urp2* could compensate for each other, at least to some extent. We thus produced double mutants from double heterozygous crosses. Zebrafish mutant for *urp1* and heterozygous for *urp2 (urp1^-/-^; urp2^-/+^*), thus lacking three out of the four copies of *urp1/2* genes, did not present any defect (Fig. 2C). In contrast, *urp1;urp2* double mutants displayed a strong spine axis curvature defect. Double mutant embryos initially developed normally, yet at 6 dpf (∼4 mm SL), a slight downward bending of the head was visible, and at the end of the second week (∼7 mm SL) the entire spine appeared bent ventrally at both ends of the body axis (Fig. 2D). At three weeks (∼8.8 mm SL), a new point of deformation could be observed just anterior to the dorsal fin resulting in an overall shape of a flat “M”. The overall shape of double mutants did not progress much during growth to adulthood and the phenotype did not seem different between males and females (Fig. 2D). The phenotype of *urp1;urp2* double mutants seemed identical to that of *uts2r3* mutants (Fig. 2E and (Zhang et al., 2018)). To characterize the spine defect of *urp1* and *urp2* mutants further, we performed alizarin stains of adult zebrafish to visualize the skeleton. This confirmed the absence of defects in *urp1* mutants (Fig. 3A-B’) as well as the kyphosis affecting the anterior half of the spine in *urp2* mutants (Fig. 3C-D’). We also found that neither *urp1* nor *urp2* mutants displayed lateral deformation (Fig3. B’’, C’’ and D’’). Further, amongst *urp2* mutants, we did not observe differences between *urp1^+/+^; urp2^-/-^* (two functional copies out of four) and *urp1^+/-^; urp2^-/-^* (only one functional copy) (Fig. 3C-D’’). *urp1;urp2* double mutants, displayed a stereotypical M shape of the spine, as described in three week-old larvae (Fig. 3 D-D’, E-E’). Of note, in contrast to what was reported in adult fish with defects in RF formation (Lu et al., 2020; Rose et al., 2020; Troutwine et al., 2020), only subtle lateral deformations were seen (Fig. 3E-E’’). Besides, we did not detect strong deformation of vertebral bones in either *uts2r3, urp1, urp2* mutants or *urp1;urp2* double mutants. Thus, our results showed that Urp1/2 signaling is required for body axis straightness during larval growth.

**Fig. 2.**
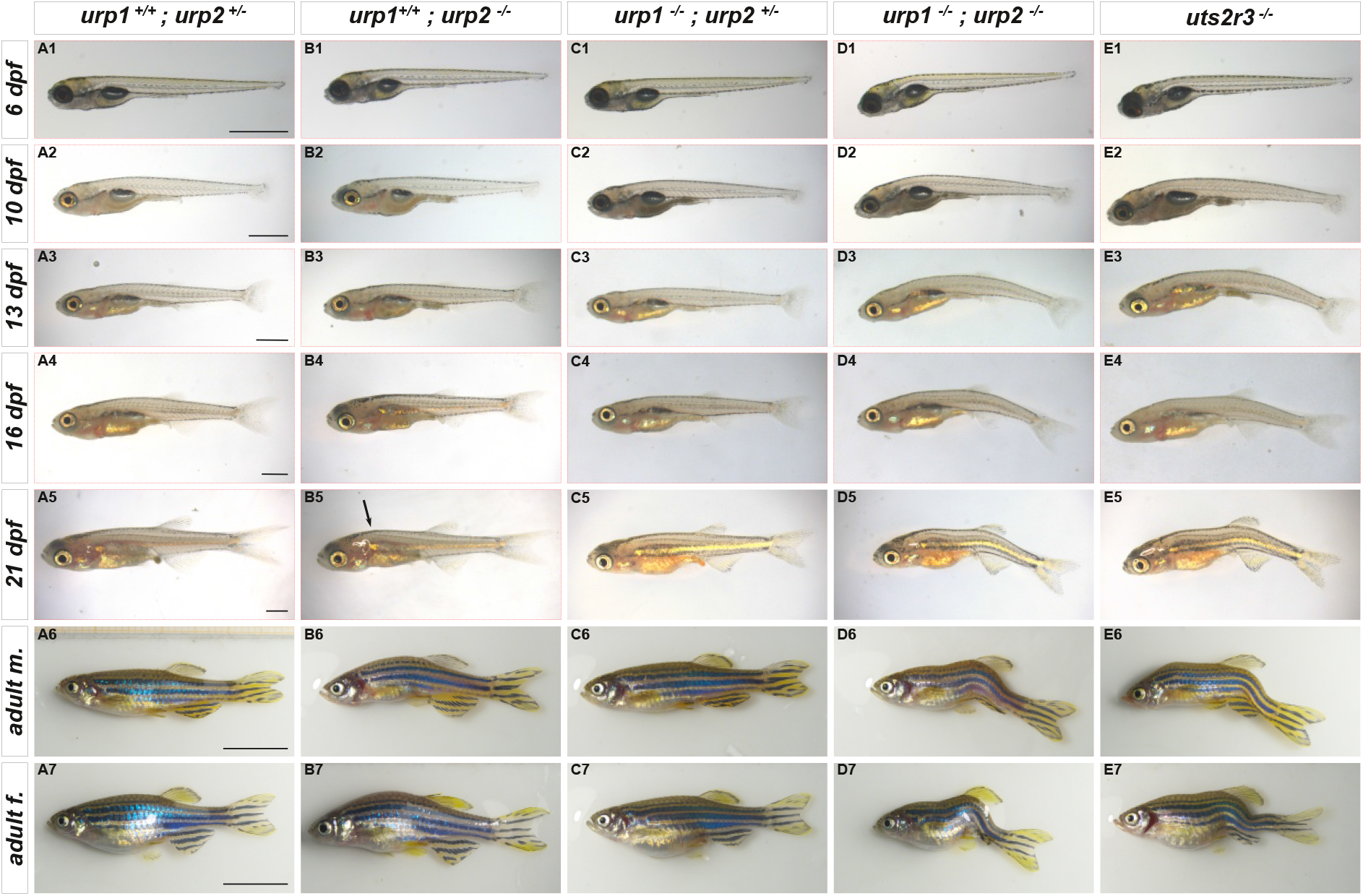
*urp2* mutants and *urp1;urp2* double mutants display progressive body axis defect. (A1-E7) Zebrafish representative of the indicated genotypes are presented at different larval stages and adults. Compared to their heterozygous siblings (A1-A7), *urp2* mutants (B1-B7) display a kyphosis which is first visible at 21 dpf. Note how the dorsal side along the head and the trunk appears as a broken line (B5, arrow. Compare to A5). Animals mutant for *urp1* and heterozygous for *urp2* do not exhibit any defect (C1-C7), but double *urp1;urp2* mutants show a strong spine axis defect visible from 6 dpf (D1-D7). This phenotype is identical to that of *uts2r3* mutants (D1-D7). For each genotype, pictures outlined in red correspond to the same animal imaged at different stages. Age – Standard length (SL) equivalence: 6 dpf = 4mm (SL); 10 dpf = 5-5.2 mm; 13 dpf = 6-6.2 mm; 16 dpf = 6.8-7.2 mm; 21 dpf = 8.5-9.2mm. Genotypes and stages as indicated. From 6 dpf to 21 dpf, scale bars represent 1mm. Adults, scale bars represent 1cm.

**Fig. 3.**
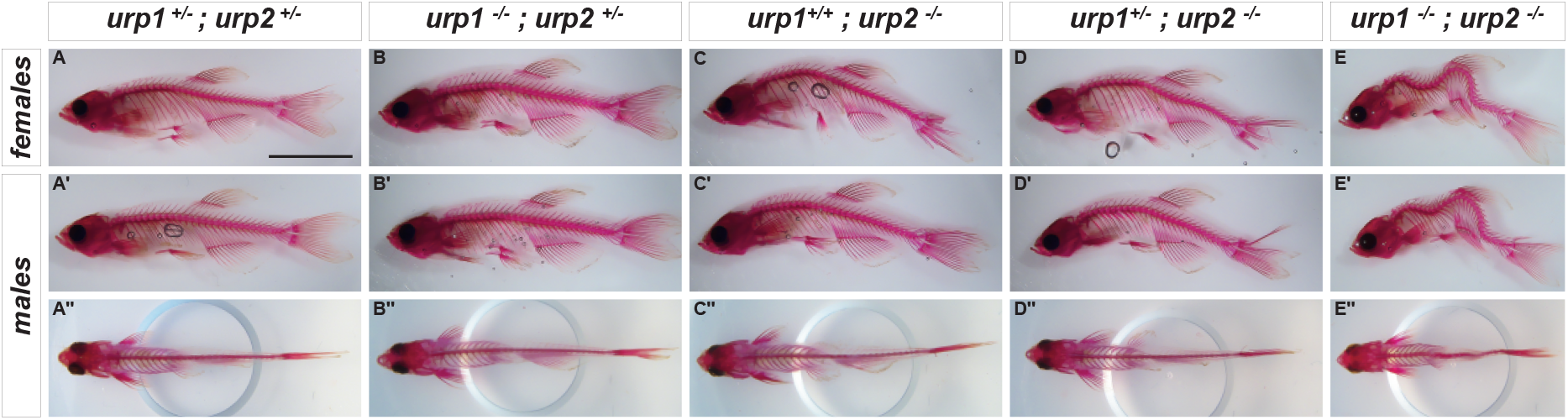
Loss of Urp2 or of Urp1 and Urp2 results in spine defect. (A – E’’) Images of alizarin stains of adult (1 year) zebrafish. Fish double-heterozygous for *urp1* and *urp2 (A-A’’),* as well as fish *urp1* mutant; *urp2* heterozygous (B-B’’) display normal phenotype. *urp2* mutants, wild-type for *urp1* (C-C’’) or heterozygous for *urp1* (D-D’’) show identical phenotypes with a strong kyphosis without lateral deformation of the spine. Double *urp1;urp2* mutants exhibit strong dorso-ventral deformation of the spine but only subtle lateral defects. A-E’, lateral views. A’’-E’’, dorsal views. Note that in dorsal view, the dorsal fin was removed to allow visualization of the spine. Scale bar represents 1 cm.

### 3.3 Osteogenesis is not impaired in *urp1;urp2* double mutants

To test if the phenotype in *urp1;urp2* double mutants could be due to congenital defect in osteogenesis, we performed calcein stain on young larvae stages (8 and 12 pdf, Fig. 4A-D) and alizarin staining in late larvae (28 pdf, Fig. 4E-F). *urp1;urp2* double mutants were compared to wild-type like *urp1^-/-^; urp2^-/+^*siblings. Strikingly, while at these stages the phenotype could be clearly seen, there was no evidence for a delay in osteogenesis nor for any congenital malformation of the vertebral bones. Thus, the phenotype observed in the absence of Urp1/2 signaling does not appear to be due to congenital defects in osteogenesis.

**Fig. 4.**
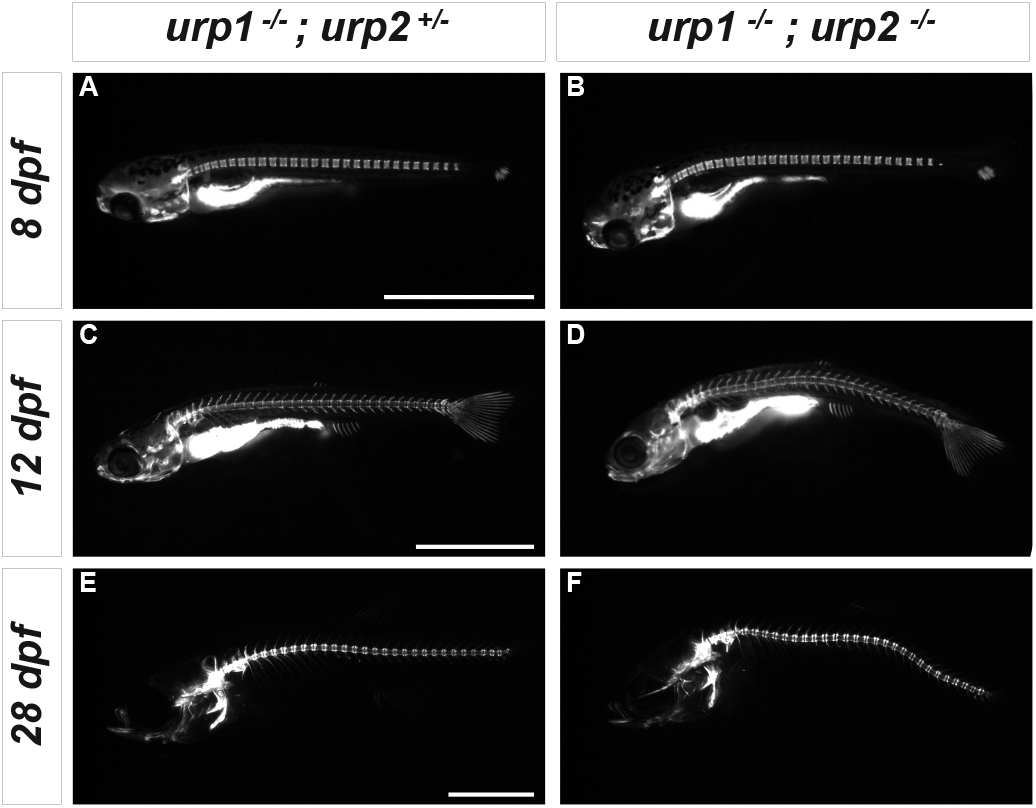
Spine defects in *urp1;urp2* double mutants are not due to abnormal osteogenesis. (A-D) Calcein stains of live *urp1^-/-^; urp2^+/-^* (A, C) and of *urp1^-/-^; urp2^-/-^* (B, D) at 8 dpf (A-B) and 12 dpf (C-D). (E-F) Alizarin stains of 28 dpf larvae. At all stages, the spine defect can be observed in *urp1;urp2* double mutants. i.e. bending of the head at 8 dpf, bending of the tail at 12 dpf and middle deformation at 28 dpf. Nevertheless, osteogenesis does not appear delayed or affected in the double mutant. Scale bars represent 2 mm.

### 3.4 Inflammation is not increased in either *urp1;urp2* or in *uts2r3* mutants

It has been reported that spine deformations in the *ptk7* and *sspo* mutants are driven by neuroinflammation (Rose et al., 2020; Van Gennip et al., 2018) and in *sspo* mutants, expression of *urp2* is impaired (Cantaut-Belarif et al., 2020; Lu et al., 2020; Rose et al., 2020). To test whether inflammation could be the cause of the deformation observed in *urp1;urp2* double mutants and *uts2r3* mutants, we used RT-qPCR to analyze the expression levels of inflammation markers in 14-day-old larvae, at the onset of the phenotype in both types of mutants. We analyzed 11 genes whose expression levels were shown to be modified in the *ptk7* mutant (Van Gennip et al., 2018). Strikingly, we did not notice any obvious increase in any of these markers in both *urp1;urp2* and *uts2r3* mutants (Fig. 5). A possible increase in the mRNA level of *irg1l*, encoding a mitochondrial enzyme involved in ROS production, was detected in *urp1;urp2* mutants but not in *uts2r3* mutants (Fig. 5), suggesting that this is not linked to the spine defect. Altogether these results suggested that the axis curvature phenotype seen in absence of Urp1/2 signaling in larvae is not caused by inflammation as a predominant defect.

**Fig. 5.**
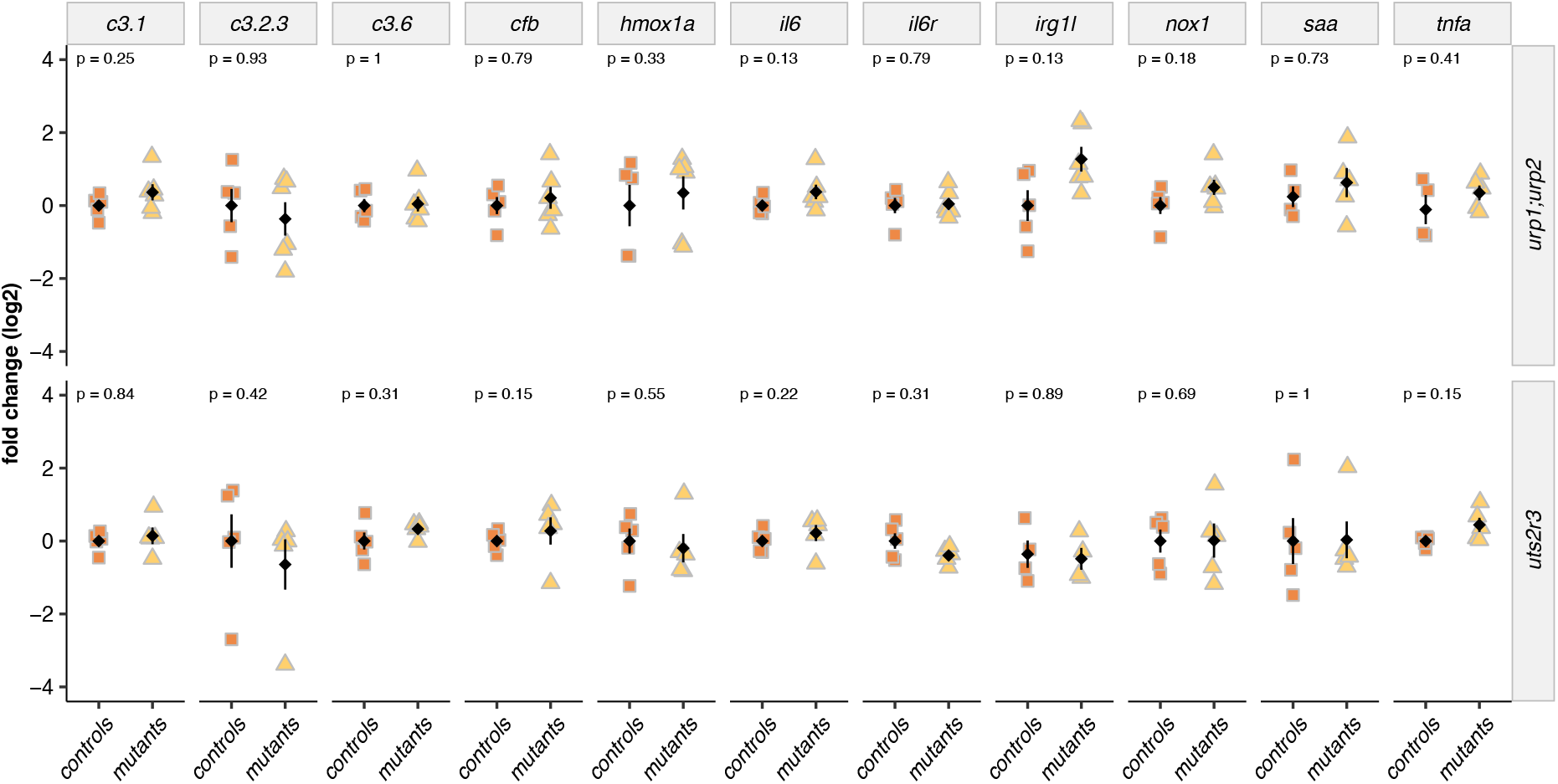
Inflammation markers are not upregulated in *urp1;urp2* mutants nor in *uts2r3* mutants. Quantitative RT-PCR analysis of mRNA levels for different inflammation markers previously shown to be associated with spine curvature (Van Gennip et al. 2018). *urp1^-/-^; urp2^-/-^* double mutants were compared to their wild-type like *urp1^-/-^; urp2^+/-^*siblings, and *uts2r3^-/-^* to their *uts2r3^+/-^* siblings. RT-qPCR were performed at 14 dpf, when the mutant phenotype starts to be clearly visible. Each point represents a pool of 5 larvae. Black diamonds and bars represent mean ± s.e.m. For each gene, p.value from Wilcoxon rank-sum tests is indicated.

### 3.5 Action of Urp1 and Urp2 is mediated by muscle contraction

Since *uts2r3* is expressed in muscles, it has been proposed that Urp1 and Urp2 could directly trigger muscle contraction (Zhang et al., 2018). This idea is supported by the observations that overexpressing either *urp1* or *urp2* in muscles leads to an upward bending of the tail in embryos and that this effect is abolished in *smoothened* mutants that lack slow-twitch muscles (Barresi et al., 2000; Lu et al., 2020). These two experiments are nonetheless not enough to demonstrate that *urp1* and *urp2* induce muscle contraction. Indeed, a bending of the tail can also be induced with Urp1 peptide injection into the CSF and the altered Hedgehog signaling in *smoothened* mutants is likely to also impact the spinal cord patterning on top of slow muscle formation. Thus, to test the idea that Urp1/2 signaling acts on muscle, we aimed at blocking muscle contractions with myosin II inhibitors (BTS and blebbistatin) and assayed the consequences on Urp2-induced tail bending in embryos. We have previously shown that a heat-shock-driven overexpression of *urp2* in embryos at 1 dpf induces an upward bending of the tail (Quan et al., 2021). Strikingly, when embryos were treated with blebbistatin prior to the heat shock, the Urp2-induced bending was abolished (Fig. 6). When embryos were treated with BTS, the upward bending was reduced but not abolished (Fig. 6). While blebbistatin can inhibit contractions of both fast and slow muscles (Eddinger et al., 2007; Limouze et al., 2004), BTS appears less potent on slow muscles (Cheung et al., 2002). Thus, these results are consistent with the fact that *uts2r3* was suggested to be expressed in slow muscles (Lu et al., 2020; Zhang et al., 2018). Altogether our results reinforced the notion that Urp1/2 signaling promotes muscle contraction to trigger the upward bending.

**Fig. 6.**
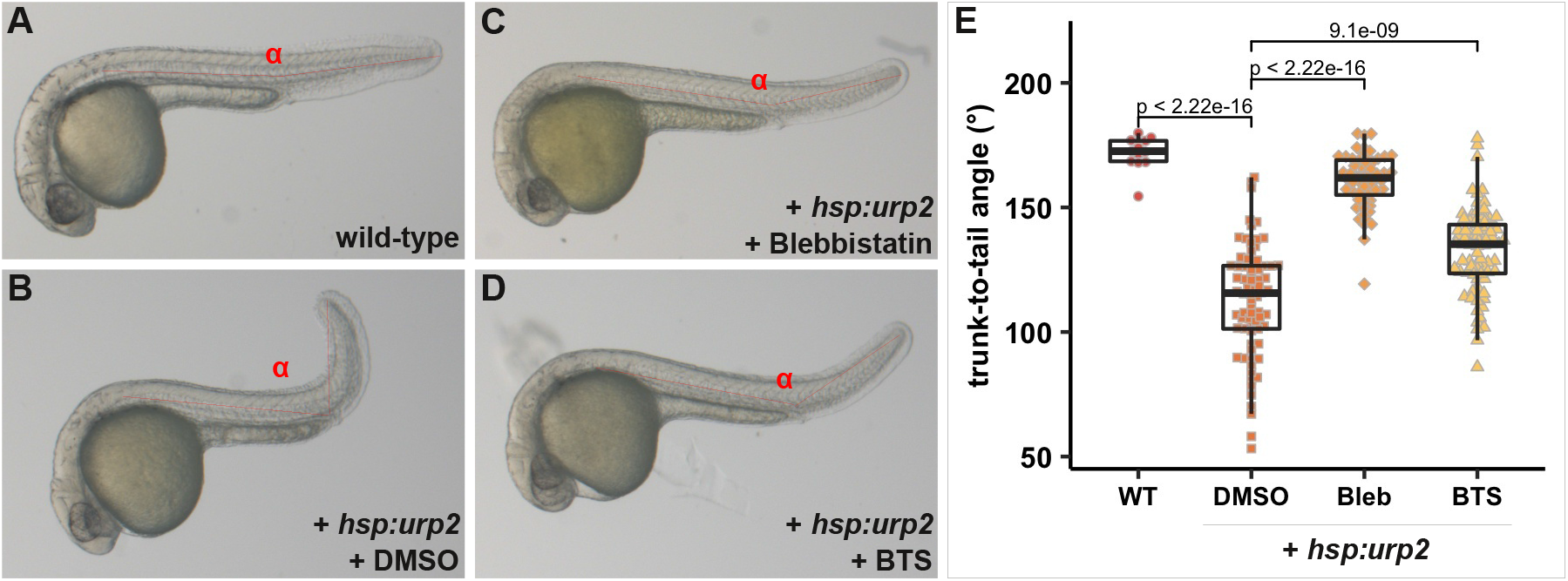
Myosin inhibitors prevent urp2-induced tail bending in embryos. Representative wild-type non-injected controls embryos (A) or wild-type embryos injected with a *hsp:urp2* transgene (B-D). 24-25 hpf embryos were placed in E3 with DMSO (B), blebbistatin (C), or BTS (D) and immediately submitted to a heatshock for 1h at 37°C and then imaged. The angle between the trunk and the tail was measured as indicated in red on each image. Data were collected from four experiments and include 13 non-injected controls, 74 embryos treated with DMSO, 58 embryos with blebbistatin and 78 embryos with BTS. p.value from pairwise Welch two sample t-test are indicated.

### 3.6 Urp signaling is functioning in larvae

While overexpressing Urp2 leads to an upward bending of the tail in embryos, most likely through dorsal muscle contraction, whether Urp1/2-Uts2r3 signaling has the same effect in larvae is not known. We thus meant to test whether Urp1/2-Uts2r3 signaling is indeed active during larval growth when the phenotype of *urp1;urp2* double mutants becomes obvious.

We have previously reported that *urp1* and *urp2* are expressed in CSF-cNs not only in embryos but also in adults (Quan et al., 2015). As for *uts2r3*, it was shown to be expressed in dorsal muscles in embryos (Zhang et al., 2018) but its expression in adults has not been described. We, therefore, used RT-qPCR to analyze the expression of *uts2r3* in various tissues in adults. Notably, we separated muscles into dorsal and ventral parts by dissecting them along the horizontal myoseptum. Consistent with what was found in embryos, our results showed that muscle expression of *uts2r3* was concentrated in dorsal fibers (Fig. 7). We also noted some expression in various tissues including the spinal cord, skin, swim bladder, and testis (Fig. 7). As for *urp1* and *urp2*, their expression was detectable primarily in the spinal cord with some expression in the brain (Fig.7), confirming our previous work (Parmentier et al., 2011; Quan et al., 2015). Of note, *urp2* appeared expressed at a much higher level than *urp1* (3-4 fold, in the spinal cord), and was also detected in the eyes (Fig. 7).

**Fig. 7.**
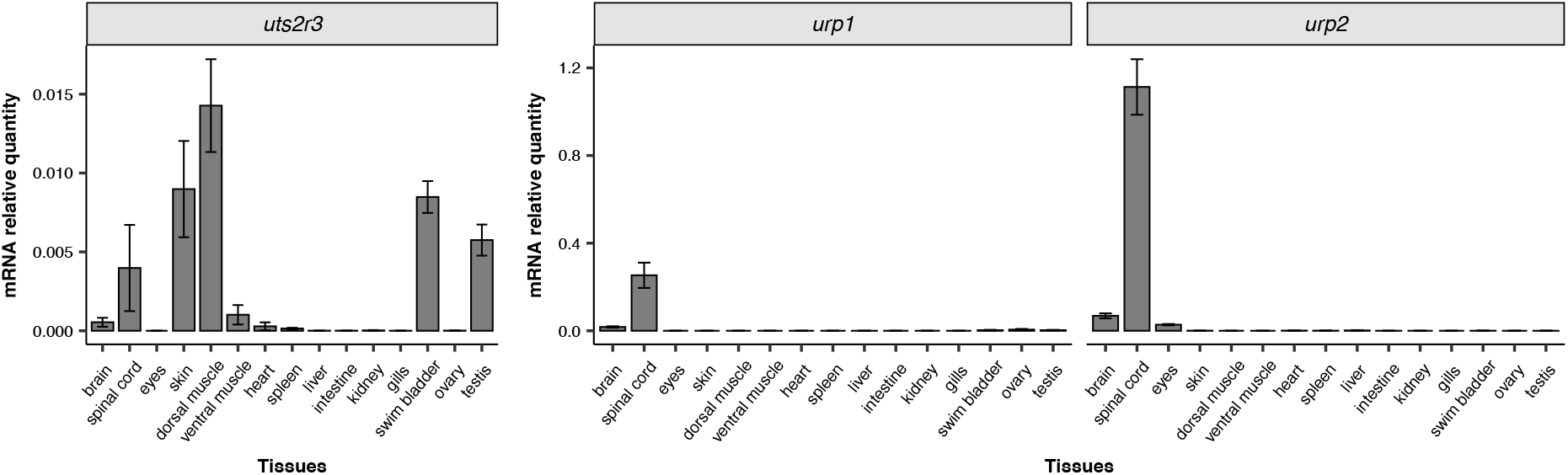
Expression profile of *uts2r3*, *urp1* and *urp2* in adult tissues. Expression levels were assayed by RT-qPCR in tissues from adult zebrafish. Expression levels are expressed as ratios to housekeeping genes. Data are means from at least three independent experiments. Error-bars are s.e.m.

In the absence of available *urp1/2* alleles that enable conditional loss-of-function to test gene function at larval stages, we aimed at inducing *urp2* overexpression in larvae. First, we injected a construct driving *urp2* under the control of the promotor of the *ubiquitin b* (ubi) gene that is active in virtually all cell types at all stages (Mosimann et al., 2011). This resulted in bending of the tail that was evident from 1 dpf and increasing with age (Fig. 8). These larvae were not raised further than 6 dpf as they could not swim nor feed properly. Nevertheless, it demonstrated that the *ubi* promotor was efficiently driving *urp2* expression and caused progressive upward bending. Next, we took advantage of the Cre/lox-system to control the timing of *urp2* overexpression in larvae. We built a transgenic construct in which the *urp2* coding sequence is separated from the *ubi* promoter by a Stop cassette, which in turn is flanked by two *loxP* sequences (*ubi:loxP-Stop-loxP_urp2;* see Methods). This construct was injected into transgenic embryos harboring a 4-hydroxytamoxifen-inducible CRE recombinase (CRE^ERt2^) driver transgene expressed in all tissues under control of the *ubi* promoter (*ubi:creERT2*) (Mosimann et al., 2011). To trigger the overexpression of *urp2*, recombination of the Stop cassette was induced by 4-hydroxytamoxifen treatment (see Methods) at 10 dpf. While, in the absence of the CRE^ERt2^-expressing transgene, 4-hydroxytamoxifen had no visible effect on zebrafish development (Fig. 9A-C’), larvae carrying and expressing CRE^ERt2^ displayed an upward bending of the tail four days after the beginning of the tamoxifen treatment (Fig. 9D-E, G-H). In some larvae, the phenotype became so strong (Fig. 9G-H, n=11/45) that they could not swim anymore and had to be euthanized. For those that could be raised to adulthood, the phenotype did not aggravate as much as in embryos, but nevertheless, the spine appeared strongly bent dorsally (Fig. 9F-F’ n=21/45). Notably, in one animal, the caudal part of the spine became bent downward, as if some compensating mechanisms allowed to replace the tail in the horizontal plane (Fig. 9I-I’). Finally, we also tried to induce *urp2* overexpression in 20 days-old larvae that carried *ubi:creERT2* and were injected with the inducible *urp2* construct. At this age, the 4-hydroxytamoxifen treatment had less effect: yet, for some animals, a slight bending of the spine could be observed a few days later (Fig. 9J-K, n=4/30). In those, the phenotype ultimately became similar to that of larvae treated at 10 dpf (Fig. 9L-L’, compare with 9F-F’). Altogether, our results suggested that Urp1/2-Uts2r3 signaling is functional during larval growth and can affect the straightness of the spine.

**Fig. 8.**
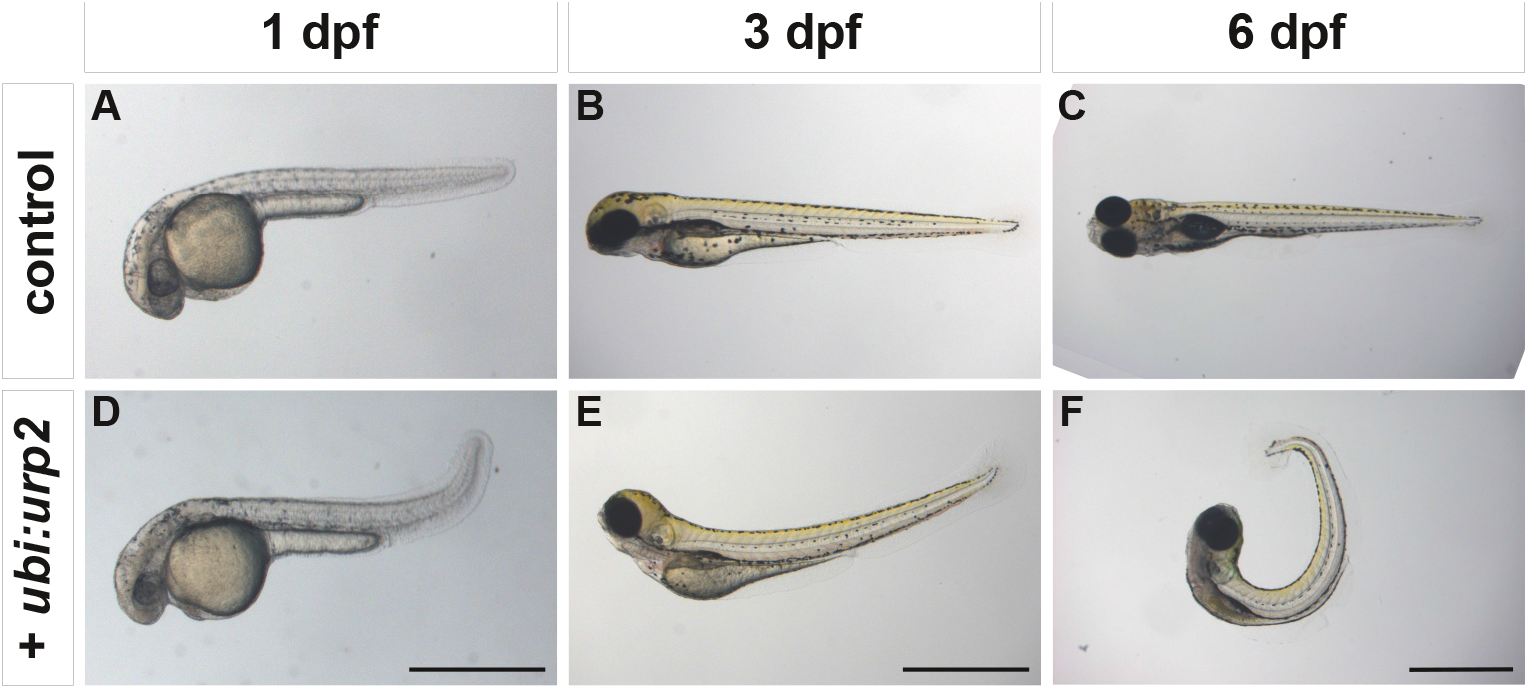
*ubi* driven overexpression of *urp2* induces an upward bending in embryos. Representative wild-type embryos, non-injected controls (A-C) or injected with an *ubi:urp2* transgene (D-F) and imaged at 1 dpf (A, D), 3 dpf (B, E) and 6 dpf (C, F). Note the progressive upward bending of the entire body axis. Scale bar: 1mm.

**Fig. 9.**
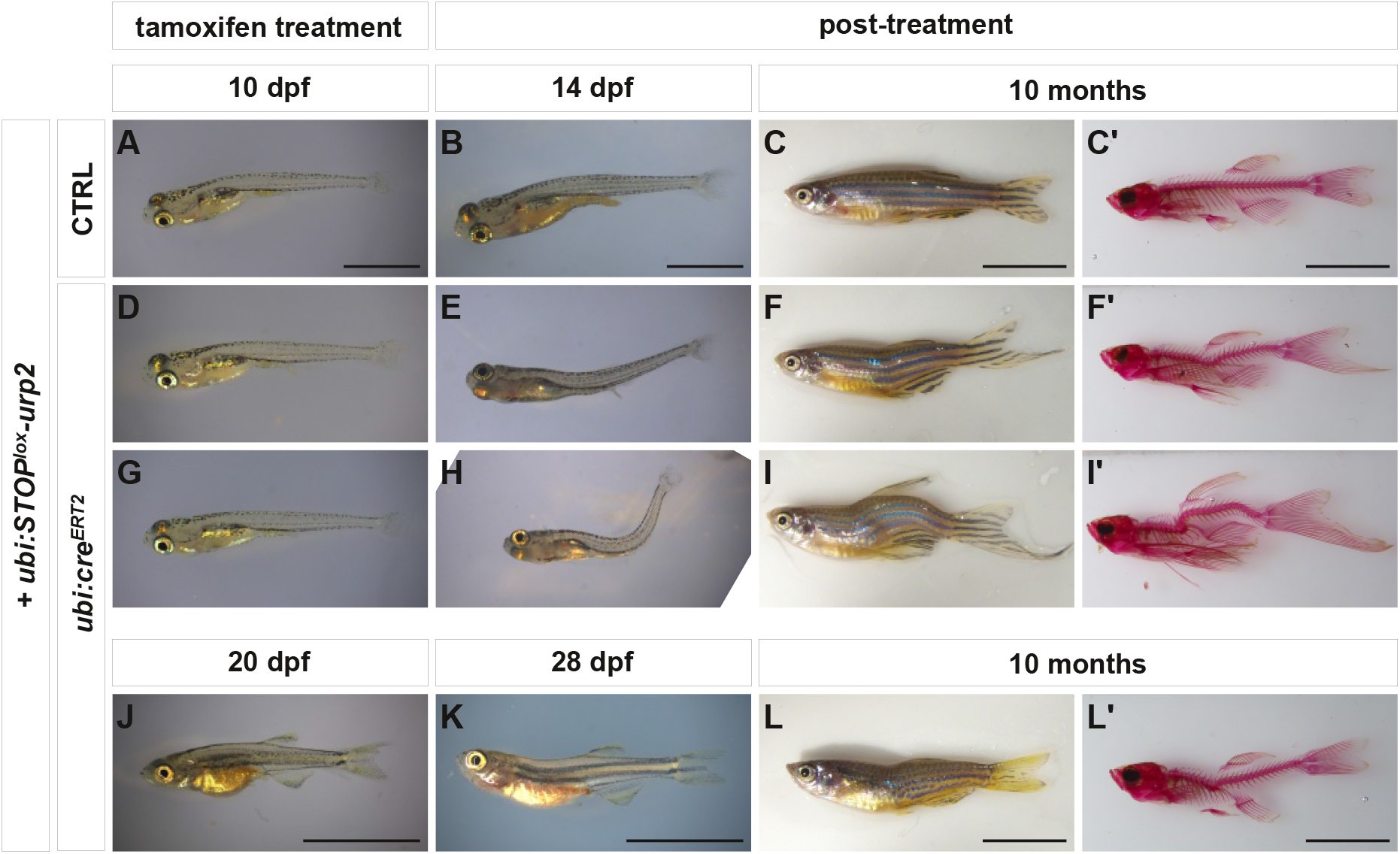
Inducible *urp2* overexpression in larvae induces spine dorsal bending. (A-C’) Control zebrafish injected with an *ubi:STOP^lox^-urp2* transgene, live (A-C) or after alizarin staining (C’). (D – L’) Transgenic *ubi:cre^ERt2^* zebrafish injected with an *ubi:stop^lox^-urp2* transgene imaged live (D-F, G-I and J-L) or after alizarin staining (F’, I’ and L’). Cre-mediated loxP recombination, which removes the stop cassette and triggers *urp2* expression, was induced with tamoxifen (4-OHT, see methods) at 10 dpf (A-I’, with two examples) or at 20 dpf (J – L’) and animals were raised until adult (10 months). Note that when treated at 10 dpf, many larvae exhibited a strong phenotype (see H) and could not be raised further. In some adults, while the spine appeared strongly bent dorsally at the level of the dorsal fin, a ventral bending was observed in the caudal region (I-I’). For each live adult zebrafish image, the adjacent image with alizarin staining corresponds to the same animal. Scale bars. 10-14 dpf: 2mm; 20-28 dpf: 5mm; adults: 1 cm.

### 3.7 Urp1/2 signaling is dispensable for embryonic development

Several studies suggested that *urp1* and/or *urp2* are required to ensure axis straightness during embryonic development (Cantaut-Belarif et al., 2020; Lu et al., 2020; Zhang et al., 2018). In particular, it was reported that morpholinos targeting *urp1* could induce a “curled-down” axis at 2 dpf. Still, none of *urp1* or *urp2* mutants nor *urp1;urp2* double mutants presented any defect during embryonic development. Likewise, *uts2r3* mutants developed normally until 6 dpf, again in contradiction with MO results (Zhang et al., 2018). A possible explanation for this discrepancy was that maternal mRNA stores could compensate for the mutations at early embryonic stages. To test for this possibility, we used RT-qPCR to analyze the expression levels of *urp1*, *urp2,* and *uts2r3* at different developmental stages. In 1-cell embryos, very low levels of *urp1* transcript could be detected but not of *urp2* nor *uts2r3*. Our analyses showed that *urp1* and *urp2* are only expressed from 24 hpf onwards and *uts2r3* from 16 hpf (Fig. 10A). Nevertheless, some protein accumulation in oocytes could explain the absence of phenotype. Despite the deformation of their spine, *urp1;urp2* double mutant females as well as *uts2r3* mutant females could lay when mated with phenotypically wild-type males. We thus crossed *urp1;urp2* double mutant females with *urp1^-/-^;urp2^-/+^* males to produce maternal and zygotic (MZ) *urp1;urp2* double mutants (50% of a clutch, the other 50% being *urp1^-/-^;urp2^-/+^*). Strikingly, these embryos did not show any defect and developed normally until 6 days just like regular zygotic double-mutants (Fig. 10C-D) Similarly, we could obtain MZ*uts2r3* mutant embryos, with no morphological defects (not shown). Altogether, this showed that the absence of embryonic phenotype is not due to compensation by stored messengers or proteins of the corresponding genes in oocytes.

**Fig. 10.**
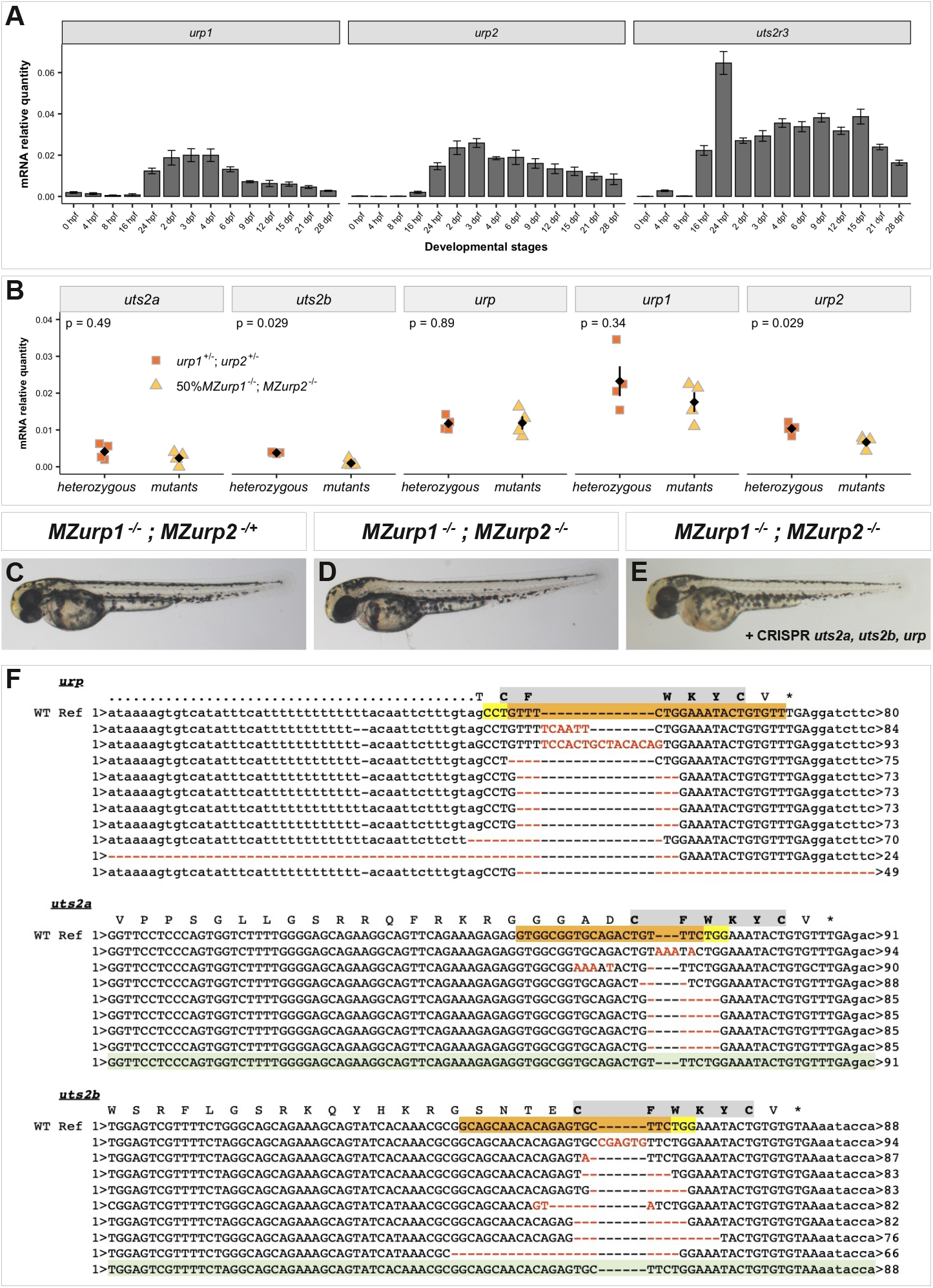
Urp1/2 signaling is dispensable for correct embryonic development. (A) Expression dynamics of *urp1*, *urp2* and *uts2r3* assayed by RT-qPCR during zebrafish development. Expression levels are expressed as ratios to housekeeping genes. Data are means from five or six independent results. Error-bars are s.e.m. (B) Expression levels of *uts2a, uts2b, urp, urp1* and *urp2* in 24 hpf embryos. Pools of embryos from clutches made of 50% MZ*urp1*^-/-^; MZ*urp2*^+/-^ (i.e. wild-type like) and 50% MZ*urp1*^-/-^; MZ*urp2*^-/-^ (ie. double mutants) are compared to 100% *urp1^+/-^;urp2^+/-^*double heterozygous. Each colored point represents a pool of 15 embryos. Expression levels are expressed as ratios to housekeeping genes. Black diamonds and bars represent mean ± s.e.m. For each gene, p.value from Wilcoxon rank-sum tests is indicated. (C-E) Representative 2-days old embryos MZ*urp1*^-/-^; MZ*urp2*^-/+^ (C), MZ*urp1*^-/-^; MZ*urp2*^-/-^ (D), and MZ*urp1*^-/-^; MZ*urp2*^-/-^ injected with three sgRNAs targeting *uts2, uts2b* and *urp* genes (E). (F) Sequences of independent clones obtained from PCR amplicons produced on DNA from a pool of 15 embryos injected with 3 sgRNA targeting *uts2, uts2b* and *urp* genes. For each gene, the reference genomic sequence, around the region encoding the core of the mature peptide, is aligned with individual sequences. The corresponding translation is indicated above the WT genomic sequence, with the core of the peptide highlighted in grey. The position of crRNA is highlighted in orange with PAM in yellow. Deleted and mutated nucleotides are in red. For both *uts2a* and *uts2b*, one non mutated sequence was obtained, highlighted in green. Overall, the observed mutation scores were: *urp* 10/10 (100%); *uts2a* 7/8 (87,5%); *uts2b* 8/9 (88,9%).

It has been reported that the absence of phenotype in some mutants is due to genetic compensation, which can be triggered by mutant mRNA decay (El-Brolosy et al., 2019). In zebrafish, there are three other genes encoding for peptides of the Urotensin 2 family. *urp* (a.k.a. *uts2d*) which is expressed in motoneurons (Quan et al., 2021, 2012), and the two *uts2* paralogues, *uts2a* and *uts2b*, expressed in neuroendocrine cells located in the caudal spinal cord, called Dahlgren cells (Parmentier et al., 2008). To test whether the absence of phenotype was due to transcriptional adaptation of either one of these three genes, we aimed to compare expression levels in double mutants to that of double heterozygous. Double mutant females were mated to wild-type males to produce clutches of 100% double heterozygous embryos, or to *urp1^-/-^;urp2^+/-^* males giving clutches with 50% of *urp1^-/-^;urp2^+/-^* and 50% of *urp1^-/-^;urp2^-/-^*(i.e., double-mutants). mRNA of 1-dpf embryos from four independent clutches were compared using RT-qPCR. Strikingly, we could not detect any increase in the expression level of *uts2a, uts2b* or *urp.* By contrast, a reduction of *urp1* and of *urp2* mRNA level was observed, most likely reflecting mRNA decay from the mutant alleles. Thus, the absence of phenotype in *urp1; urp2* double mutant embryos did not seem to be due to genetic compensation by genes encoding other members of the Urotensin 2 family.

Nevertheless, to confirm further the absence of a requirement of Urp1/2 signaling during embryogenesis, we aimed at inhibiting all five *uts2/urp* genes. To do so, we used CRISPR-Cas9 to inhibit *uts2a, uts2b* and *urp* in MZ*urp1;urp2* double mutants. The core of all Urotensin 2 related peptides is a cyclic hexapeptide (CFWKYC) conserved in all species examined so far in vertebrates (Vaudry et al., 2015), and any mutation in this sequence should result in an inactive peptide. Thus, we designed gRNAs targeting the coding sequence of the core of each three peptides and injected simultaneously the three guides in 50% MZ*urp1;urp2* clutches. Sequence analyzes of targeted regions showed high levels of mutations (Fig. 10F) suggesting efficient inactivation of the three targeted genes. Yet, these *uts2a;uts2b;urp* triple crispant – *urp1;urp2* double MZmutant embryos, although completely deprived of Uts2/Urps peptides, never displayed the typical curled-down phenotype reported with morpholino (Fig. 10E). Altogether, these results strongly suggested that Urp1/2 signaling is dispensable for correct embryonic development.

### 3.8 The Pkd2 pathway controls embryonic axis straightness independently of URP1-2 signaling

While overexpression of Urp1/2 can affect axis straightness in embryos, the fact that Urp1/2 signaling appeared dispensable during embryonic development suggested the existence of Urp1/2-independent pathway(s) that control axis straightness. To test this possibility, we turned to the *pkd2* mutant that displays an upward bending of the tail (curled-up phenotype) at 2 dpf (Schottenfeld et al., 2007). We reasoned that if this phenotype depended upon Urp1/2 – Uts2r3 signaling, it should be abolished in an Uts2r3-deficient background. We thus used CRISPR-Cas9 to inhibit *pkd2* function, with two gRNAs targeting the *pkd2* gene, in 50% MZ*uts2r3^-/-^* clutches. Injected embryos were imaged at 2 dpf to measure the angle between the trunk and the tail and then genotyped. Strikingly, while CRISPR-Cas9 -mediated mutagenesis of *pkd2* largely reproduced the upward bending of the tail observed in the *pkd2* mutant, the absence of Uts2r3 did not affect the severity of the phenotype (Fig. 11). This result is consistent with the notion that during embryogenesis, at least one Urp1/2-independent pathway, dependent on Pkd2, is involved in axis straightness control.

**Fig. 11.**
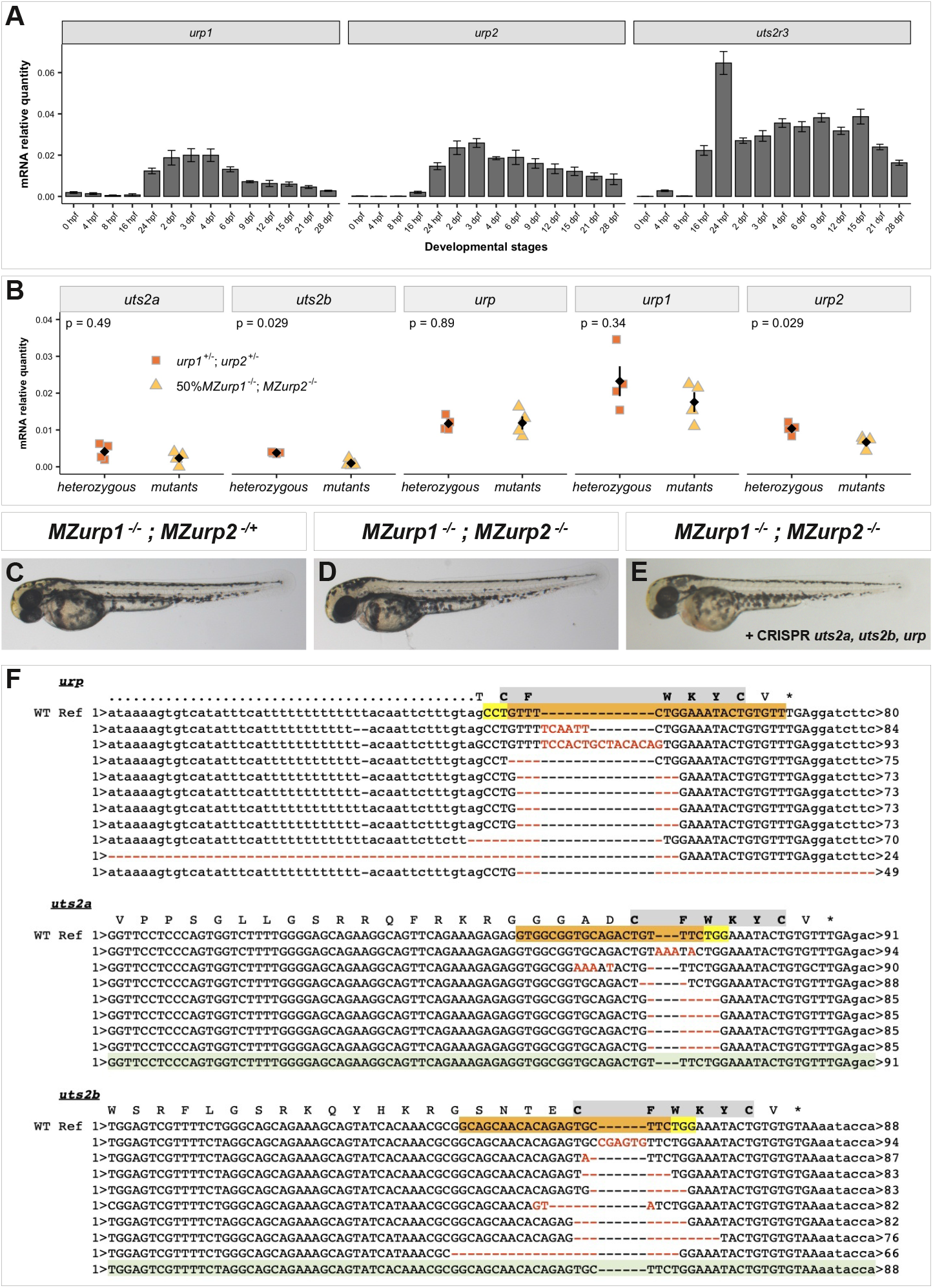
Pkd2 curly-up phenotype does not depend on Uts2r3. Females *uts2r3^-/-^*mutant were mated with *uts2r3^+/-^* males, to produce clutches made of 50% MZ*uts2r3^+/-^* and 50% MZ*uts2r3^-/-^*. Embryos were injected at 1-2-cell stage with two sgRNAs targeting the *pkd2* gene. At 2 dpf, embryos were images and then genotyped. MZ*uts2r3^+/-^* (A) and MZ*uts2r3^-/-^* (B) non-injected controls do not display any abnormal phenotype. By contrast, MZ*uts2r3^+/-^* (C,E) and MZ*uts2r3^-/-^* (D,F) crispants for *pkd2* exhibit mild (C-D) to severe (E-F) curly-up phenotype. (G). Quantification of the measured angles between the trunk and the tail revealed no difference between the two genotypes. Data were collected from two experiments and include 39 MZ*uts2r3^+/-^*and 40 MZ*uts2r3^-/-^*. The angles were measured on images of embryos without knowing the genotype. p.value from Welch two sample t-test is indicated.

## 4 Discussion

Since seminal studies have shown that zebrafish can be used as a model to study idiopathic scoliosis (Buchan et al., 2014; Gorman and Breden, 2009; Hayes et al., 2014), a condition affecting between 1% and 4% of adolescents (Cheng et al., 2015), considerable progress have been made toward understanding the mechanisms involved in spine and body axis formation in vertebrates (Bagnat and Gray, 2020; Muñoz-Montecinos et al., 2021). Urp1 and Urp2, two neuropeptides of the Urotensin 2 family, have been recently shown to be involved in this process (Cantaut-Belarif et al., 2020; Lu et al., 2020; Wang et al., 2020; Zhang et al., 2018). By analyzing zebrafish mutant for *urp1* or *urp2* and *urp1;urp2* double mutants, we show here that, while being dispensable for embryogenesis, these two neuropeptides act together during larval growth to promote correct vertebral body axis. The redundancy between these two peptides is consistent with the fact that they are largely co-expressed in the same cells and were found to be equipotent to activate a human Uts2r in a cell assay (Quan et al., 2015). We also show that the Urp1/2 effect depends on myosin II, reinforcing the idea that their action relies on the modulation of muscle tone.

Urp1/2 are produced by the CSF-cNs, under the control of signals from the Reissner fiber whose formation requires cilia (Cantaut-Belarif et al., 2020; Lu et al., 2020; Rose et al., 2020). Previous results in both cilia-defective and *sspo* mutants showed that when such animals can be raised to adulthood they exhibit strong spine deformation along both dorso-ventral axis and medio-lateral axis (Hayes et al., 2014; Lu et al., 2020; Rose et al., 2020; Troutwine et al., 2020). By contrast, *urp1;urp2* double mutants and *uts2r3* mutants have strong deformations along the dorso-ventral axis but only subtle deformations along the medio-lateral axis. Interestingly, ventral CSF-cNs, in which Urp1/2 are produced, were suggested to respond to spine contractions along the dorso-ventral axis (Hubbard et al., 2016), while dorsal CSF-cNs seem involved in response to lateral bending (Böhm et al., 2016). Nevertheless, both types of CSF-cNs require the Reissner fiber to detect spinal curvature (Orts-Del’Immagine et al., 2020). Thus, during larval growth, it could be hypothesized that cilia and the Reissner fiber are required for the regulation of at least two signals to control the vertebral body axis: one acting on ventral CSF-cNs, regulating spine straightness along the dorso-ventral axis through Urp1/2 signaling; and a second one, still unknown, acting on dorsal CSF-cNs and controlling body shape along the medio-lateral axis.

Another difference between cilia-defective and *sspo* mutants, and *urp1;urp2* double mutants and *uts2r3* mutants is that inflammatory signals were proposed to be the main cause of the spine deformations observed in the former (Rose et al., 2020; Van Gennip et al., 2018) but that we did not observe an increase of these signals in the latter. Even if we have only analyzed inflammatory signals in 14 dpf larvae and cannot rule out a possible increased in inflammation at latter stages, our results show that inflammation is not the primary cause of the deformation of the spine in absence of Urp1/2 signaling, in contrast to cilia-defective and *sspo* mutants. Interestingly, anti-inflammatory treatment could ameliorate the phenotype of *sspo* mutant embryos but did not affect the observed downregulation of *urp2* (Rose et al., 2020). The cause of inflammation in absence of Reissner fiber remains unknown and it should be noted that another study did not reveal such an increase in inflammatory signals in absence of Reissner fiber (Cantaut-Belarif et al., 2020). Also, the mechanisms of action of inflammation on spine morphogenesis, as well as whether there is a link between inflammation and *urp1/2* regulation, remain to be studied, but a possibility could be that inflammation prevents normal activity of both ventral and dorsal CSF-cNs, thus participating to the overall impairment of spine morphogenesis observed in absence of Reissner fiber.

Urp1/2 are thought to act through the receptor *uts2r3* whose mutant exhibits the same phenotype as *urp1;urp2* double mutants (Zhang et al. 2018 and our results). Also, Uts2r3 is required for the upward bending of the tail induced by *urp1/2* overexpression. Since *uts2r3* is expressed in dorsal muscles both in embryos (Zhang et al., 2018) and in adults (our work) and since we found that the effect of Urp1/2 can be blocked by myosin II inhibitors, these peptides most likely function by regulating muscle tone. It was reported that the effect of *urp1/2* overexpression in embryos is not blocked by alpha-bungarotoxin (Zhang et al., 2018), an inhibitor of acetylcholine receptor, suggesting that Urp1/2 signaling does not rely on classical neuromuscular signal and supporting the idea of a direct effect on muscles through Uts2r3. Interestingly, Urp, another peptide of the Urotensin2 family, is expressed in motoneurons (Quan et al., 2021) and would have also been an ideal candidate to activate Uts2r3. Yet, we have previously found that the loss of function of *urp* has no visible effect on zebrafish development and that Urp is not required for Urp1/2-induced upward bending of the tail (Quan et al., 2021). If this suggests that Urp1/2 signals directly on muscles through Uts2r3, then the question remaining is how. CSF-cNs have different types of projections but mostly toward different types of neurons in the spinal cord (Djenoune et al., 2017; Wu et al., 2021). Some projections toward dorsal muscles were described (Wang et al., 2020) but whether they actually reach muscles is not clear. One possibility could be that Urp1/2 are secreted in the CSF or outside the spinal cord and diffuse or are transported by the bloodstream toward muscles. In support of this idea, in embryos, the injection of Urp1 peptides into the CSF can induce an upward bending of the tail (Zhang et al., 2018) and the related Uts2 neuropeptide have been reported to exert multiple endocrine effects (Vaudry et al., 2015). Alternatively, it should also be noted that we detected expression of *uts2r3* in the spinal cord. It is thus possible that the action of Urp1/2 relies on a second population of neurons, expressing *uts2r3*, that in turn signals on muscle. Experiments based on tissue specific inactivation of *uts2r3,* for example using spatially controlled expression of Cas9 (Donato et al., 2016), should allow to demonstrating in which cell type Uts2r3 is required.

Previous work on the function of Urp1/2, focusing on embryogenesis, reported that the injection of morpholinos targeting *urp1* or *uts2r3* resulted in a curled-down phenotype at 1-2 dpf (Zhang et al., 2018). By contrast, we found that mutants for *uts2r3, urp1, urp2* and *urp1;urp2* double mutants exhibit a wild-type phenotype during embryogenesis (i.e. up to 5 days), even in the absence of maternal protein and mRNA stores. This suggests that Urp1/2-Uts2r3 signaling is actually dispensable for embryogenesis. Consistent with this idea, we have previously reported that, in Xenopus, CRISPR-mediated inhibition of *utr4* (the counterpart of zebrafish *uts2r3)* does not provoke axis curvature defect in embryos but results in curled-down phenotype in tadpoles, similar to that observed in zebrafish *uts2r3* mutant larvae (Alejevski et al., 2021). This suggests that the mechanisms involved in body axis morphogenesis are conserved between fishes and amphibians. Discrepancies between mutant and morphant phenotypes are not rare (Kok et al., 2015). On some occasions they can be explained by genetic compensation (also referred as transcriptional adaptation) which can be triggered by nonsense-mediated mRNA decay (El-Brolosy et al., 2019; Ma et al., 2019). Our results suggest that this is not the case here. So, could the phenotype of *urp1* and *uts2r3* morphants be not specific? Morpholinos are known to frequently induce non-specific effects. In particular, they can provoke toxicity due to p53 activation, which can lead to body axis defects (Bedell et al., 2011). This phenomenon can be tested by co-injecting a morpholino targeting p53, but this experiment was not reported for *urp1* nor *uts2r3* morphants (Zhang et al., 2018). The most classic approach to address the specificity of a phenotype in a morphant is to perform an RNA rescue experiment (Eisen and Smith, 2008). This was done by Zhang et al. (2018) for *urp1* morphant, but in this particular case, the experiment is not really conclusive. Indeed, injection of *urp1* mRNA was also reported to rescue the phenotype of *zmind* and *sspo* mutants (Cantaut-Belarif et al., 2020; Zhang et al., 2018). If *urp1* mRNA can rescue mutants for other genes than *urp1* itself, then it cannot be used to demonstrate that the phenotype *urp1* morphant is specific to *urp1* knock-down. This difficulty should not arise with *ust2r3,* but, sadly, Zhang et al. did not show an mRNA rescue experiment for morpholinos targeting this gene. Thus, at this stage, it is not possible to tell whether the phenotypes of *urp1* and *uts2r3* morphant embryos are specific or not.

Nevertheless, several other studies suggest that Urp1/2-Uts2r3 signaling is functional during embryogenesis. On the one hand, overexpressing Urp1 or Urp2 by different means in 1 dpf embryos results in a backward bending of the tail (Cantaut-Belarif et al., 2020; Lu et al., 2020; Quan et al., 2021; Zhang et al., 2018). On the other hand, the expression of *urp1* and/or *urp2* is reduced in *zmind* and *sspo* mutants and injection of *urp1* mRNA can rescue their phenotypes (Cantaut-Belarif et al., 2020; Lu et al., 2020; Zhang et al., 2018). Downstream of the Reissner fiber, monoamines seem involved in the regulation of *urp1/2* expression (Cantaut-Belarif et al., 2020; Wang et al., 2020; Zhang et al., 2018). Altogether, these results converge to a model in which Urp1/2-Uts2r3 signaling could be involved in body axis formation during embryogenesis. If so, why is there no embryonic defect in mutants for *uts2r3, urp1, urp2* or *urp1;urp2* double mutants, even when produced by homozygous mutant females? The most reasonable explanation seems to be that at least one other independent mechanism, acts in parallel with Urp1/2 downstream of the Reissner fiber. This hypothesis is supported by several lines of evidence. First, in *sspo* mutant embryos, the expression of *urp2* was found to be reduced, but not that of *urp1* (Cantaut-Belarif et al., 2020; Rose et al., 2020). Yet, neither *urp2* mutants (this work) nor morphants (Zhang et al., 2018) display defective embryonic axis curvature. Also, we found that the curled-up phenotype of *pkd2* mutants is not affected by the absence of Uts2r3, showing that Pkd2 is required for an Urp1/2 independent mechanism regulating the morphogenesis of the embryonic axis. Further investigations will be needed to fully delineate the contribution of Urp1/2 during embryogenesis.

## Acknowledgments

We thank Céline Maurice, Philippe Durand and Jean-Paul Chaumeil (PhyMA MNHN) for zebrafish care. We are grateful to Jean-Paul Concordet (StrInG, MNHN) for his advices on CRISPR experiments; Christine Vesque and Pierre-Luc Bardet (IBPS, SU) for critical reading of the manuscript and Sylvie Schneider-Maunoury and Isabelle Anselme (IBPS, SU) for helpful discussions.

## Contributions

Conceptualization: Hervé Tostivint, Guillaume Pézeron

Methodology: Hervé Tostivint, Guillaume Pézeron

Investigation: Feng B. Quan (initial production of *urp1* and *urp2* mutants), Teddy Mohamad (Calcein staining and myosin inhibitor assay), Anne De Cian (sgRNA and Cas9 protein production), Christian Mosimann (ubi:STOPflox construct), Anne-Laure Gaillard, Guillaume Pézeron

Formal analysis: Anne-Laure Gaillard, Guillaume Pézeron

Validation: Anne-Laure Gaillard, Guillaume Pézeron

Visualization: Guillaume Pézeron

Writing – original draft: Guillaume Pézeron

Funding acquisition: Hervé Tostivint, Guillaume Pézeron

## KEY RESOURCES TABLE

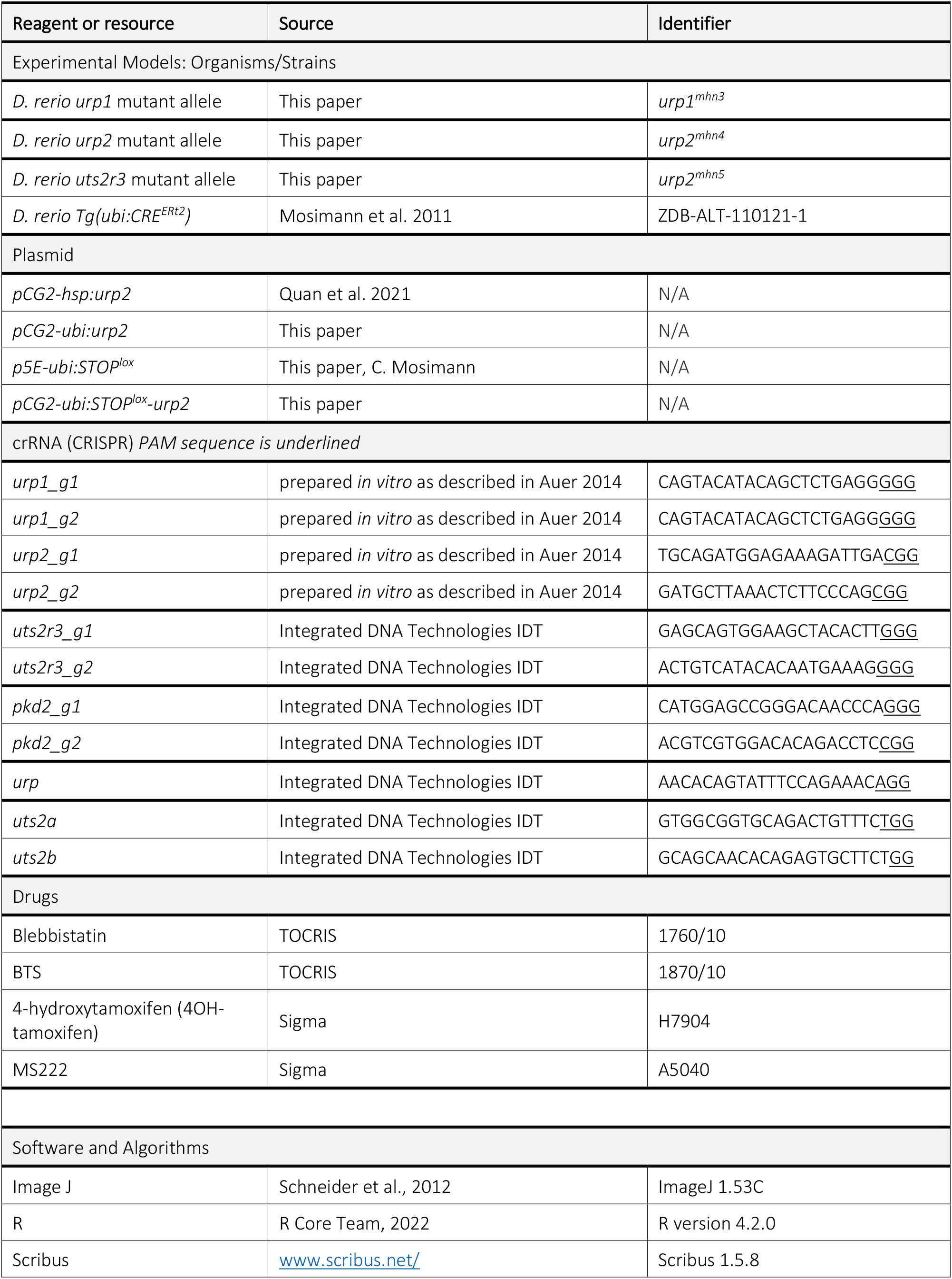

---

**Table S1.**
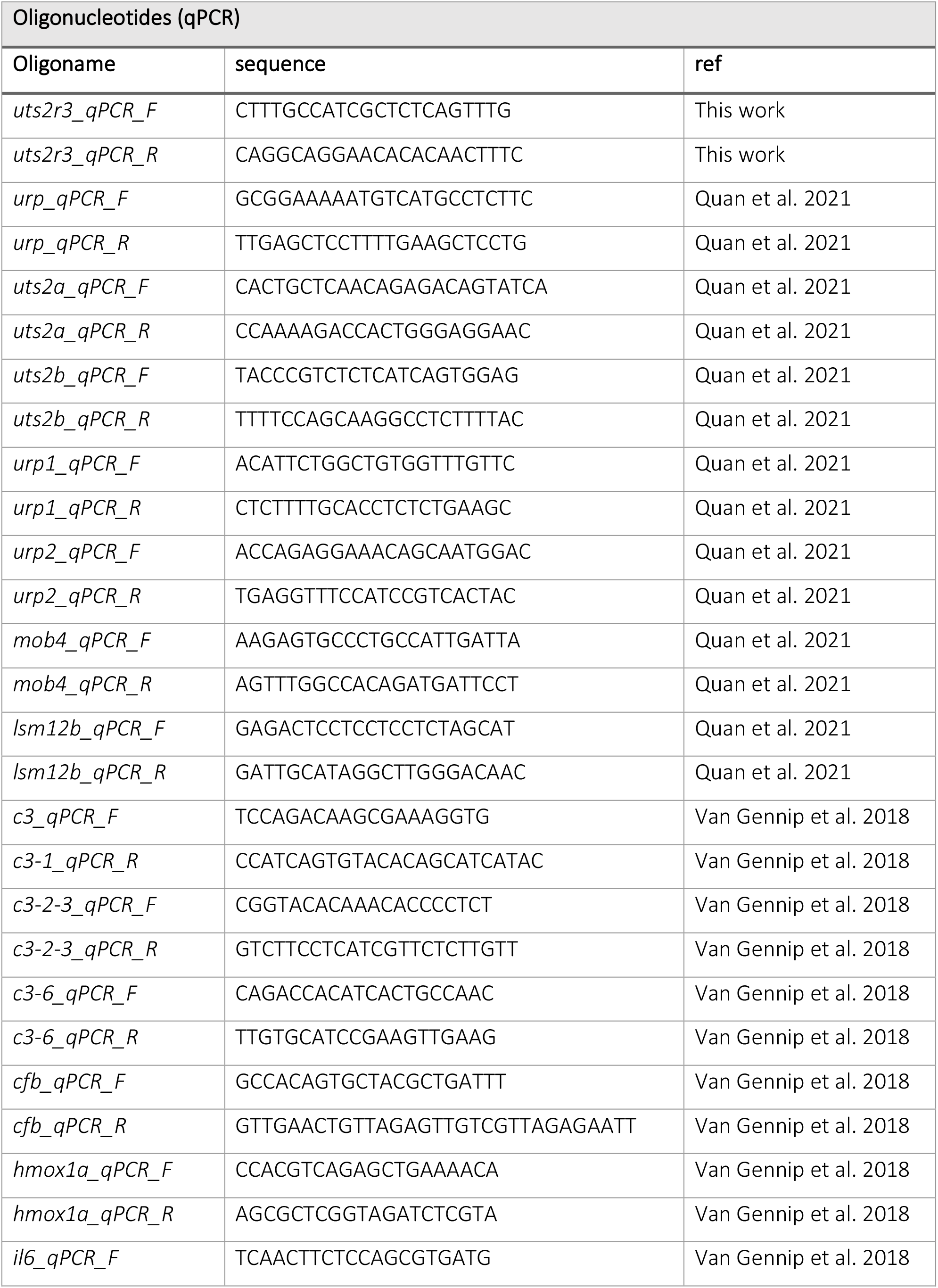
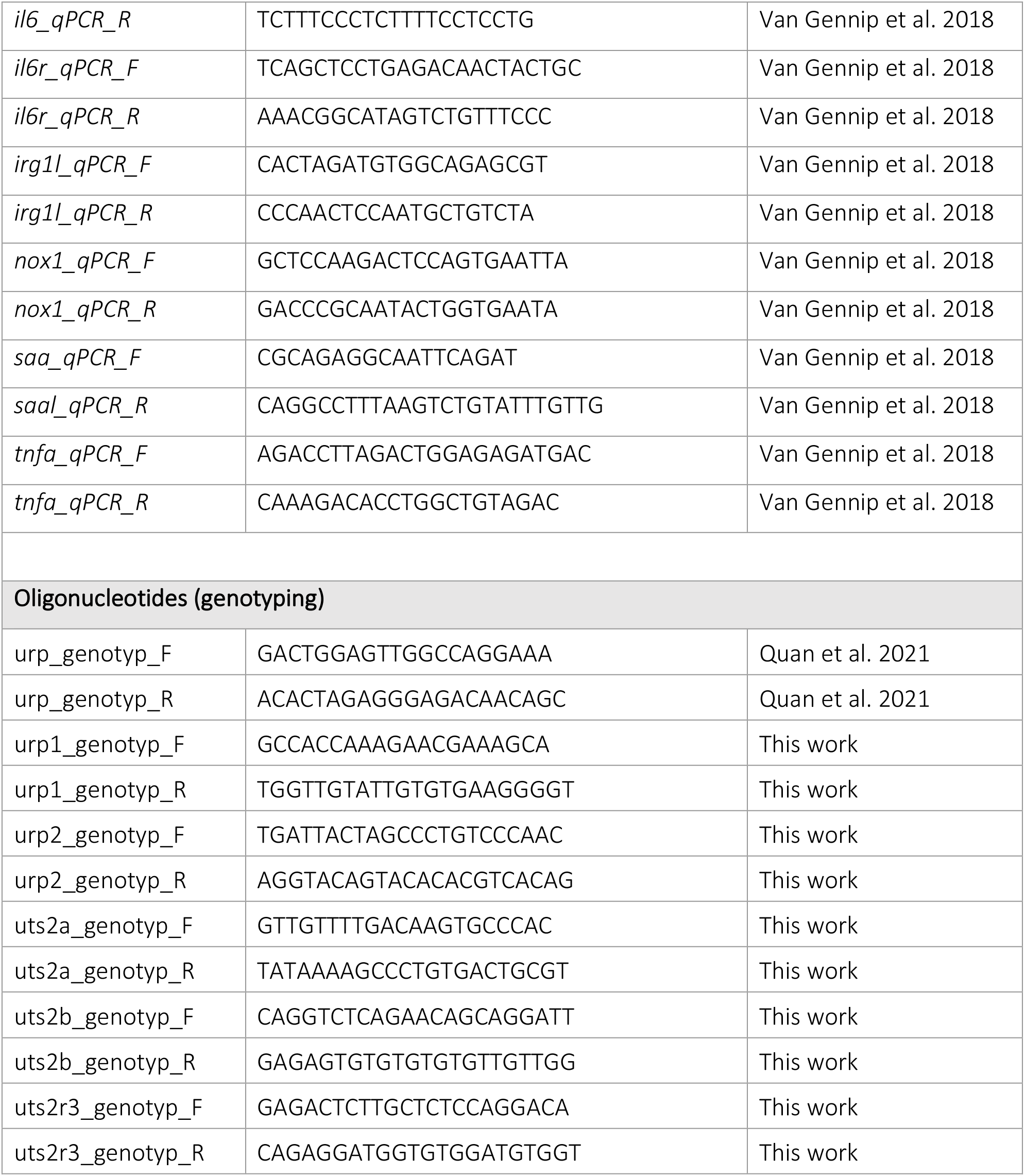

